# The influence of reproductive stage on cerebellar network connectivity across adulthood

**DOI:** 10.1101/2021.10.06.462014

**Authors:** Hannah K. Ballard, T. Bryan Jackson, Tracey H. Hicks, Jessica A. Bernard

## Abstract

Sex-specific differences in the aging cerebellum may be related to hormone changes with menopause. We evaluated the influence of reproductive stage on lobular cerebellar network connectivity using data from the Cambridge Centre for Ageing and Neuroscience repository. We used raw structural and resting state neuroimaging data and information regarding age, sex, and menopause-related variables. Crus I and II and Lobules V and VI were our cerebellar seeds of interest. We characterized reproductive stage using the Stages of Reproductive Aging Workshop criteria. Results show that postmenopausal females have lower cerebello-striatal and cerebello-cortical connectivity, particularly in frontal regions, along with lower connectivity within the cerebellum, compared to reproductive females. Postmenopausal females also exhibit greater connectivity in some brain areas as well. Differences begin to emerge across transitional stages of menopause. Further, results reveal sex-specific differences in connectivity between female reproductive groups and age-matched male control groups. This suggests that menopause may influence cerebellar network connectivity in aging females, and sex differences in the aging brain may be related to this biological process.

**Highlights:**

- Lobular analysis of cerebellar network connectivity across reproductive stages
- Postmenopausal females show lower cerebellar connectivity, compared to reproductive
- Connectivity differences begin to emerge in transitional stages of menopause
- Age-matched male control groups show distinct patterns of cerebellar connectivity
- Reproductive stage influences cerebellar network connectivity in aging females

## 1. Introduction

By 2060, the number of American adults age 65 and over is expected to nearly double in size (Administration on Aging, 2020). This will have a significant financial impact on society as healthcare needs rise. Further, the burden of functional declines with age reduces quality of life for older adults. Along with normative age-related declines, one in nine older adults age 65 and over suffer from Alzheimer’s disease (AD) (The Alzheimer’s Association, 2021). Thus, the hardships of this age-related disease, in addition to the challenges experienced in healthy aging, have major repercussions for aging Americans and society more broadly. As such, a thorough understanding of the factors that contribute to functional declines and differences with age is important for improving quality of life in the aging population, as well as reducing societal costs. As the brain is impacted in both healthy aging (Hausman et al., 2020; Jernigan et al., 2001; Raz et al., 2010, 2005; Walhovd et al., 2011; Woodruff-Pak et al., 2010) and AD (Braak and Braak, 1991; Henneman et al., 2009; Leal and Yassa, 2013; Tabatabaei-Jafari et al., 2017), uncovering the neural underpinnings of age-related functional changes and differences will provide key new information for combating cognitive and motor declines in later life.

Though aging research traditionally focuses on age-related differences in the cerebral cortex (Du et al., 2006; Golomb et al., 1993; Greenwood, 2000; Mander et al., 2014, 2013; Seidler et al., 2010; Tisserand and Jolles, 2003), the cerebellum (CBLM) is also considerably impacted by aging both cross-sectionally and longitudinally (Bernard et al., 2015, 2013, Bernard and Seidler, 2014, 2013; Han et al., 2020; Raz et al., 2005; Tabatabaei-Jafari et al., 2017; Woodruff-Pak et al., 2010). The CBLM has smaller volume in advanced age (Jernigan et al., 2001; Raz et al., 2013, 2010, 2005, 2001) and has shown earlier senescence than cortical regions more commonly associated with aging (Woodruff-Pak et al., 2010). Further, the CBLM has a sizable role in age-related behavioral differences; work suggests that smaller CBLM volumes in advanced age are associated with slower gait and balance difficulty (Bernard and Seidler, 2013; Rosano et al., 2007), as well as cognitive and intellectual performance differences (Bernard et al., 2015; Lee et al., 2005; Paul et al., 2009). Importantly, CBLM atrophy has been linked to the clinical progression of AD (Tabatabaei-Jafari et al., 2017), further highlighting the importance of understanding the CBLM in adulthood. In addition to volumetric differences, functional connectivity in CBLM networks is impacted in advanced age. Relative to young adults, older adults show disrupted resting state connectivity in CBLM-cortical networks and asynchronous connectivity in CBLM-subcortical networks (Bernard et al., 2013; Hausman et al., 2020).

Notably, females are disproportionately impacted by age-related functional challenges, relative to males. Older adult females show larger deficits in postural control and balance, higher rates of injury from falls, and steeper declines in global cognition than their male counterparts (Hartholt et al., 2011; Kim et al., 2010; Panzer et al., 1995; Proust-Lima et al., 2008; Stevens and Sogolow, 2005). Interestingly, several investigations note sex-specific differences in the CBLM with age. Females consistently have smaller CBLM volumes than males (Rhyu et al., 1999; Ruigrok et al., 2014), even when controlling for height (Raz et al., 2001, 1998). Relatedly, females are more critically affected by AD; both the prevalence and severity of symptoms is higher in females than males (Carter et al., 2012; Mazure and Swendsen, 2016). In fact, roughly two-thirds of those that suffer from AD are females (Hebert et al., 2013). Moreover, rates of CBLM atrophy with AD are greater in females (Tabatabaei-Jafari et al., 2017). Together, the literature suggests that females endure more severe consequences with aging, making it exceedingly important to understand the underlying mechanisms. Thus, exploring biological differences between sexes may help explain the differential impact of aging on females.

Specifically, menopause may contribute to sex differences in aging with respect to both brain and behavioral measures, and later life outcomes. Menopause and the associated changes in sex hormones have been linked to declines in cognition in older adult females (Gurvich et al., 2018; LeBlanc et al., 2001; Weber et al., 2012). Work by Weber et al. (2014, 2013) demonstrates that postmenopausal females show worsened performance on delayed verbal memory and phonemic fluency tasks, as well as deficits in verbal learning and motor function, compared to reproductive and perimenopausal females. Menopause is marked by the cessation of ovarian function in older adulthood, which is accompanied by a decrease in estrogen levels, and research suggests that estrogen may operate as a cognitive protective factor (Fischer et al., 2014). In aging females, estrogens have been shown to prevent neurodegeneration and, in turn, cognitive decline (Daniel, 2013). This neuroprotective characteristic of estrogen is largely supported by work investigating hormone therapy. Hormone therapy improves cognitive function (Duka et al., 2000) and is related to decreased risk of dementia and AD in older females (Gurvich et al., 2018; LeBlanc et al., 2001; Li and Singh, 2014). Complementing this idea, endogenous hormone fluctuations during the menstrual cycle have been associated with changes in resting state connectivity networks (Petersen et al., 2014; Pletzer et al., 2016). Thus, as hormonal fluctuations modify intrinsic connectivity in networks related to cognition and aging, hormone changes with menopause may contribute to differences in cognition as well as brain networks in advanced age. As menopause results in a significant decrease in estrogen levels, this biological factor, specific to females, may partially underlie the exaggerated impact of aging on females.

Chiefly, the CBLM is susceptible to the effects of estrogen. Estrogen receptors are present in the CBLM (McEwen, 2002) and local estrogen synthesis regulates CBLM neurotransmission (Hedges et al., 2018, 2012). Postmenopausal females undergoing estrogen therapy demonstrate greater grey matter volumes in the CBLM compared to those not undergoing treatment (Boccardi et al., 2006). In addition, both postmenopausal females undergoing estrogen therapy and young females in the reproductive stage have displayed larger grey matter concentrations in the CBLM compared to postmenopausal females not receiving hormone therapy (Robertson et al., 2009). Thus, the use of hormone therapy for natural age-related decreases in estrogen with menopause brings postmenopausal females to a comparable level with that of young females, in terms of CBLM volume. The benefits of estrogen therapy on CBLM morphology have also been correlated with improvements in cognition (Ghidoni et al., 2006). Given the impact of estrogen on CBLM physiology and function in postmenopausal females, along with the resulting effects on cognition, it is important to consider the relationship between menopause-induced hormone changes and modifications in the CBLM with older age.

Collectively, the literature suggests that the CBLM may be a key region impacted by menopause-induced hormone fluctuations, which has wide-ranging functional implications. However, despite extensive work on aging, CBLM connectivity in the context of menopause has not been sufficiently studied. Further, research investigating age effects on the CBLM generally adopts a gross anatomical approach (Jernigan et al., 2001; Raz et al., 1998; Rhyu et al., 1999; Rosano et al., 2007; Tabatabaei-Jafari et al., 2017), focusing on the entire CBLM or large subregions, rather than using specific lobules to drive analyses. Though this work is informative for elucidating gross CBLM organization and function, it is important to include lobular analyses for a more detailed perspective on the CBLM in advanced age. Work by Bernard et al. (2013) underscores the importance of investigating the aging CBLM with a regional approach, as specific lobules are differentially impacted in younger and older adults. To address these gaps, we examined resting state connectivity in CBLM networks across female reproductive stages, using a lobular approach. Then, we looked at relative male comparisons using age-matched male control groups to evaluate sex-specific differences in CBLM network connectivity. This work advances our understanding of the divergent functional trajectories of aging females and males, by investigating a relatively understudied biological factor with a unique analytical approach.

## 2. Methods

### 2.1. Participants

For the current study, we used raw imaging data from the Cambridge Centre for Ageing and Neuroscience (Cam-CAN) repository (Shafto et al., 2014; Taylor et al., 2017), which was collected from a cross-sectional, lifespan sample of healthy adults. We acquired structural and resting state magnetic resonance imaging (MRI) data and information regarding age, sex, and menopause-related variables for 652 subjects. To account for potential differences in brain organization (Levy and Reid, 1978; Li et al., 2014), we first excluded left-handed individuals (n = 52), as determined by the Edinburgh Handedness Inventory (Oldfield, 1971). Any individual with a negative score, indicating left-handedness, was excluded; thus, all right-handed individuals were included in our sample. A handedness score was not available for two subjects, and as such, those individuals were also excluded. Further, four subjects did not have resting state MRI data and were excluded from analyses. One additional subject was excluded for poor data quality, as their data contained significant motion artifacts that could not be effectively corrected during image processing. Lastly, three subjects were excluded as less than 40 volumes of resting state data were acquired/available for these subjects, though the Cam-CAN imaging parameters call for a total of 261 volumes. Thus, before performing age-matching, which resulted in additional exclusions, our initial sample for the current study included 590 righthanded subjects (297 females), ages 18-88.

### 2.2. Reproductive Stage

Female reproductive stage was categorized using the Stages of Reproductive Aging Workshop Criteria, as outlined by Harlow et al. (2012). Females were divided into four stage groups: reproductive, perimenopausal, early postmenopausal, and late postmenopausal. To assign females to these groups, we used data from the home interview portion of the Cam-CAN study, which included information regarding the length of menstrual cycle in days, age when menstruation stopped, and current age of subjects. To distinguish between reproductive and perimenopausal females, we used the length of menstrual cycles. Females that reported menstrual cycle lengths of less than 60 days were classified as reproductive, and females with cycles that spanned 60 days or more were determined to be in perimenopause. To accurately separate postmenopausal females into early and late groups, we used age when menstruation stopped and current age to calculate the time, in years, since the onset of menopause. Those that had undergone menopause within the last 9 years were classified as early postmenopausal, and those that had been postmenopausal for 9 years or more were assigned to the late group.

Females without data regarding these menopause-related variables were classified based on age (n = 36). As the average age of onset for menopause is roughly 49 years (Palmer et al., 2003; Schoenaker et al., 2014; Wang et al., 2018), the following age cut-offs were used to assign females lacking menstrual data to reproductive, perimenopausal, early postmenopausal, and late postmenopausal groups, respectively: 18-39, 40-49, 55-70, and 71+. Females age 50-54 who did not have enough information to classify reproductive stage based on menstrual data were excluded from final analyses (n = 5), due to considerable variability in reproductive stage for this age range (Kato et al., 1998; Morabia and Costanza, 1998; Palmer et al., 2003).

To better understand whether the results from our female reproductive stage comparisons are a product of menopause-induced hormone changes, rather than solely a factor of age, we also formed male control groups by performing 1:1 age-matching between sexes. We then mirrored the female comparisons with these male groups. Each female subject was matched to a single male subject using age, as well as two quality assurance variables (number of outlier scans and maximum motion) produced as second-level covariates during image processing. Notably, females and males do not significantly differ in either number of outlier scans (*t*(587.99) = −1.40, *p* = 0.16) or maximum motion (*t*(577.17) = −1.68, *p* = 0.09), thus data quality is comparable between sexes. Where there were several males of the same age, the subject with the number of outlier scans closest to that of the female in question was chosen as a match. Further, where the number of outlier scans was identical between potential male matches, the male with the closest value for maximum motion observed during scanning was matched to the female. As such, males and females were matched based on both age and data quality, where necessary. Males that were not matched to a female were excluded from final analyses (n = 72). The same approach was used in instances where there were more females than males of a particular age. Thus, females that did not receive a valid male match were also excluded from final analyses (n = 71). This approach allows us to compare between sexes using groups of equal age makeups and sample sizes, helping account, at least in part, for the inherent relationship between age and menopause.

The sample size and average age for each female reproductive group and each male control group, after performing age-matching, is presented in **Table 1**. After applying age-matching to both sexes to achieve equal age-makeups and samples sizes, the final sample used in our analyses consisted of 442 subjects (221 females), ages 18-87 (mean age 55.48 ± 18.71). Importantly, there is notable heterogeneity in age across female reproductive stages, even between reproductive and late postmenopausal groups (**Figure 1**). This overlap in age between female groups further suggests that we are not looking solely at age impacts but at a more complex process, that being the menopausal transition over the course of the female lifespan.

**Table 1.** Group characteristics. Sample size, average age, and standard deviation of age for each female reproductive group and male control group after applying 1: 1 age-matching. Due to age-matching, the resulting sample size and age of female and male groups are equal. Therefore, the numbers presented here correspond to both sexes.

| Stage | Sample Size | Age |
| --- | --- | --- |
| Reproductive | 72 | 36.22 ± 8.96 |
| Perimenopause | 25 | 40.28 ± 9.14 |
| Early Postmenopause | 26 | 56.96 ± 5.14 |
| Late Postmenopause | 98 | 73.11 ± 7.74 |

**Figure 1.**
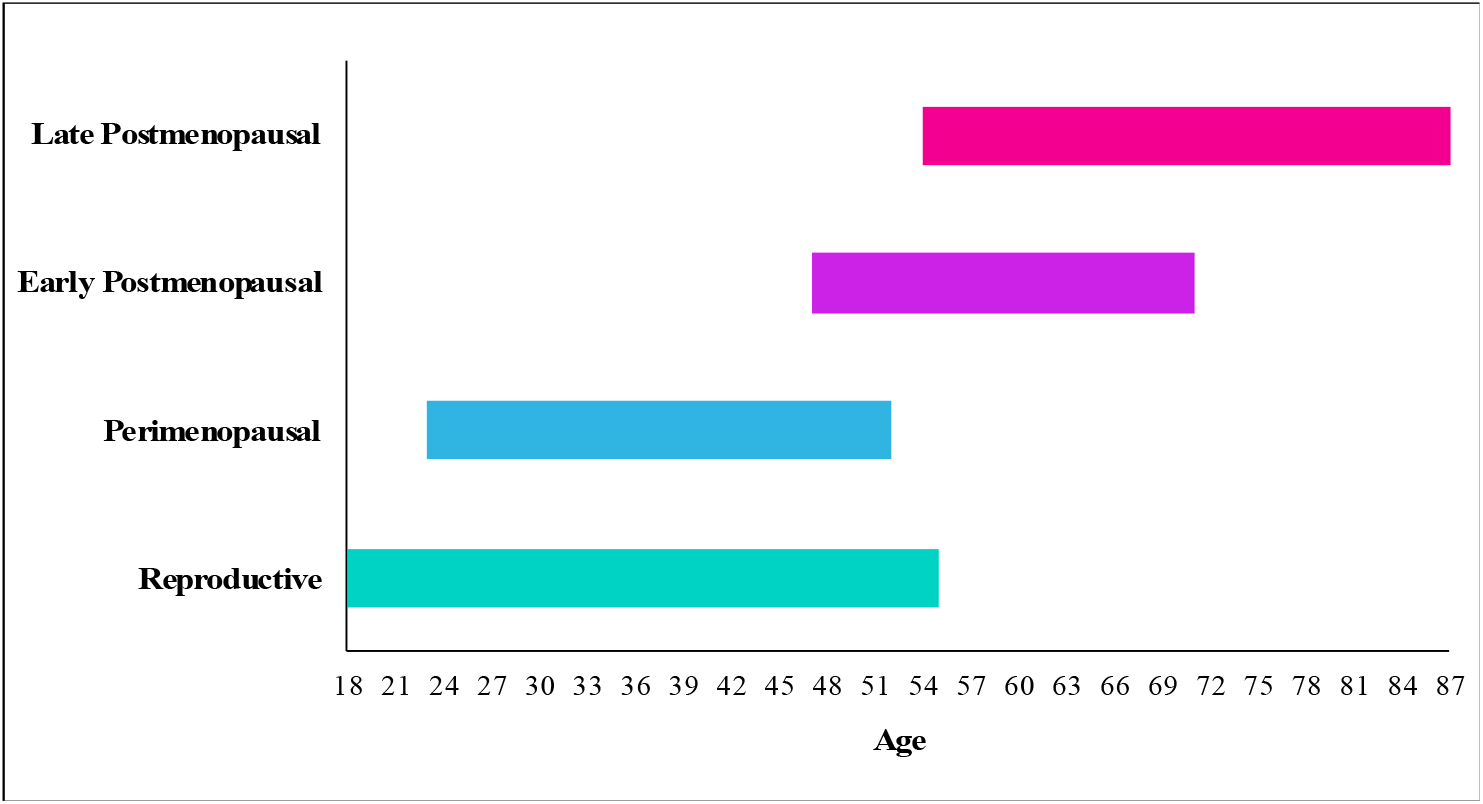
Age distribution of female reproductive stages. Age range for each female group was calculated after applying age-matching exclusions. Reproductive: n = 72, age range = 18-55; Perimenopausal: n = 25, age range = 23-52; Early Postmenopausal: n = 26, age range = 47-71; Late Postmenopausal: n = 98, age range = 54-87.

### 2.3. Connectivity Analyses

Imaging data from the Cam-CAN repository were collected with a 3T Siemens TimTrio, using a voxel size of 3 × 3 × 4.4 mm, acquisition time of 8.5 minutes, and TR (repetition time) of 1.97 seconds, for resting state scans. We used the raw T1 MPRAGE structural scans and raw resting state EPI scans for our analyses. A detailed overview of the data collection parameters and sample characteristics is available in Taylor et al. (2017) and Shafto et al. (2014). For the current study, we used Crus I, Crus II, Lobule V, and Lobule VI of the right hemisphere for our CBLM seeds, as defined by the SUIT atlas (Diedrichsen, 2006; Diedrichsen et al., 2009) and used in our past work (Bernard et al., 2016, 2013, 2012). Given that our analyses were performed on a right-handed sample, corresponding to dominance in the right CBLM hemisphere, we localized our seeds of interest to this specific hemisphere. These particular seeds were chosen because they show age differences in cortical and subcortical connectivity (Bernard et al., 2013; Hausman et al., 2020), as well as volume (Bernard et al., 2015; Bernard and Seidler, 2013), making them especially pertinent to our interests here. Further, we were interested in the impact of reproductive stage on connectivity in both cognitive (Crus I & II) and motor (Lobule V & VI) components of the CBLM (Stoodley and Schmahmann, 2009).

Image processing and connectivity analyses were conducted in version 19b of the CONN toolbox (Whitfield-Gabrieli and Nieto-Castanon, 2012), paralleling methods used in prior work (Bernard et al., 2017; Hausman et al., 2020). The default preprocessing pipeline was used to perform functional realignment and unwarping, along with subject motion estimation and correction, functional and structural centering to (0, 0, 0) coordinates, functional slice-timing correction, functional outlier detection using the Artifact Rejection Toolbox (ART) with a conservative 95^th^ percentile threshold, functional and structural segmentation of grey matter, white matter, and cerebrospinal fluid, as well as normalization to Montreal Neurological Institute (MNI) space, and spatial smoothing using a Gaussian kernel of 5 mm full width at half-maximum (FWHM). Data were denoised with a band-pass filter of 0.008-0.099 Hz, and the global-signal z-value threshold was set at 3 with a 0.5 mm threshold for subject motion correction. 6-axis motion information and frame-wise outliers, de-spiked during denoising to conform to the global mean, were included as first-level covariates. The frame-wise time series was averaged across subjects, and 6-axis motion covariates were formed by averaging absolute values of each axes’ average.

First-level analyses were conducted to quantify individual seed-to-voxel relationships, computed using a whole-brain bivariate correlation approach, for all subjects. Then, we looked across age-matched subjects with reproductive stage for females and male control groups as a second-level covariate. Second-level analyses were primarily focused on the female groups in order to investigate the impact of reproductive stage on CBLM connectivity within females. In addition, second-level analyses were conducted across age-matched male control groups in order to qualitatively compare connectivity patterns between sexes. All group-level comparisons were carried out using a voxel threshold of p < 0.001 and an FDR-corrected cluster threshold of p < 0.05. Imaging analyses were conducted using the advanced computing resources provided by the Texas A&M High Performance Research Computing organization.

## 3. Results

### 3.1. Female Reproductive Stage Comparisons

Compared to reproductive females, late postmenopausal females show lower CBLM-cortical connectivity, particularly in frontal regions, across all CBLM seeds (**Figure 2**; **Supplementary Table 1**). Along with several frontal areas, connectivity in late postmenopausal females was lower in parietal, temporal and occipital cortices. In addition, late postmenopausal females exhibit lower CBLM-subcortical connectivity, especially in the basal ganglia, across seeds. Lower connectivity is observed in the caudate, putamen, amygdala, cingulate gyrus, and insular cortex. Further, with each seed, lower connectivity within the CBLM is observed in late postmenopausal females, relative to reproductive.

**Figure 2.**
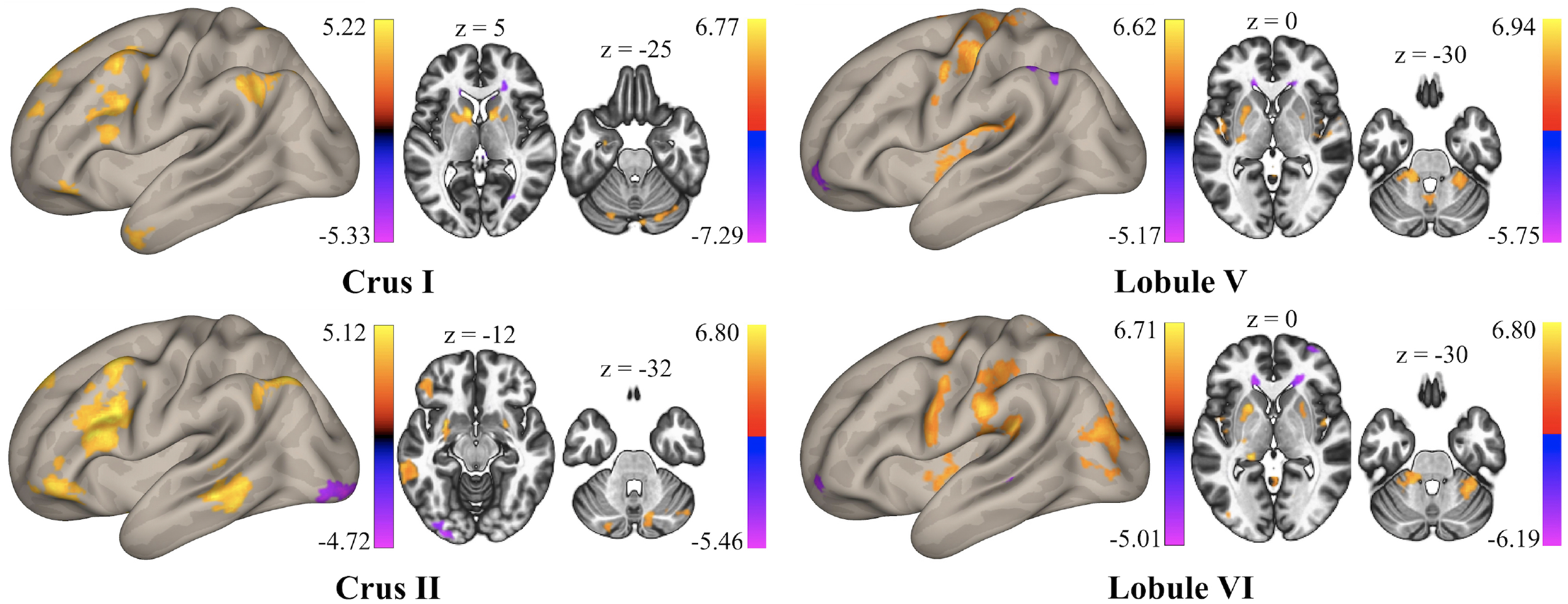
Reproductive females > late postmenopausal females. Seed-to-voxel contrasts between reproductive and late postmenopausal females for Crus I & II (left) and Lobule V & VI (right). Orange/yellow indicates areas of greater connectivity in reproductive females, and blue/purple indicates greater connectivity in late postmenopausal females.

In contrast, late postmenopausal females also demonstrate greater connectivity between CBLM seeds and some cortical areas. For instance, greater connectivity is observed in the middle occipital gyrus along with parietal regions, such as the precuneus cortex and angular gyrus. Interestingly, late postmenopausal females exhibit greater connectivity in a few subcortical regions as well, including the corona radiata, a component of the basal ganglia, and the thalamic radiation, which connects the prefrontal cortex to the thalamus. Notably, late postmenopausal females do not show greater connectivity within the CBLM, for most seeds.

Importantly, however, most of the significant differences for this particular comparison indicate greater connectivity in reproductive females, compared to late postmenopausal females. In fact, 64% of all significant clusters correspond to greater connectivity in reproductive females. This indicates that CBLM resting state connectivity is generally lower in late postmenopausal females, relative to reproductive, though areas with greater connectivity are present as well.

Notably, differences in CBLM connectivity begin to emerge across transitional stages of menopause (perimenopause and early postmenopause). For instance, when comparing reproductive to early postmenopausal females, the majority of differences indicate lower connectivity in early postmenopausal females in areas such as the amygdala, hippocampus, putamen, precuneus, and frontal lobe (**Figure 3A**; **Supplementary Table 2**). However, no significant differences appear between reproductive and perimenopausal females. Between perimenopausal and late postmenopausal females, all associations signify lower connectivity in late postmenopausal females, primarily in frontal and striatal areas (**Figure 3B**; **Supplementary Table 3**). On the other hand, only one significant difference is present between perimenopausal and early postmenopausal females; perimenopausal females exhibit greater connectivity between Crus I and the precuneus (*cluster size* = 91; *x* = 12, *y* = −40, *z* = 40; *P*_(*FDR*)_ = 0.013; *B* = 0.180). Finally, when comparing early and late postmenopausal females, we see a fairly equal distribution of differences in both cortical and subcortical locations (**Figure 3C**; **Supplementary Table 4**), suggesting that there are some instances where connectivity is higher in both groups.

**Figure 3.**
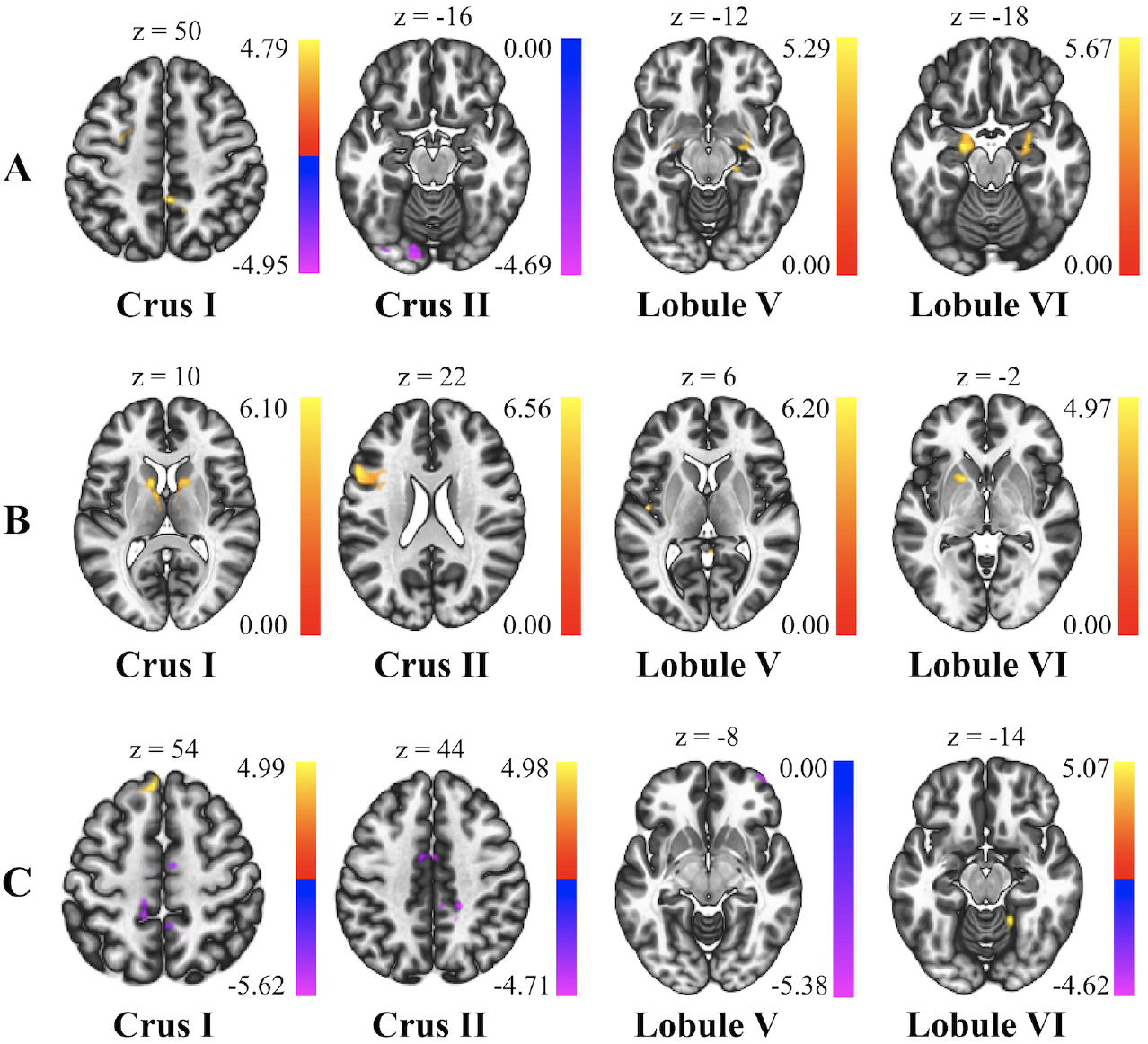
Comparisons with transitional female stages. Largest clusters, based on cluster size, for the following comparisons: A: reproductive females > early postmenopausal females, B: perimenopausal females > late postmenopausal females, and C: early postmenopausal females > late postmenopausal females. Color bars represent connectivity in the primary group (left of >), relative to the secondary (right of >), on a scale of lower connectivity (blue/purple) to greater connectivity (orange/yellow).

### 3.2. Age-Matched Male Control Comparisons

While age-matched male control groups show some of the same cortical and subcortical differences as relative female groups, there are also several distinct differences in the observed patterns (**Figure 4**, **Supplementary Table 5**). Similar to late postmenopausal females, male controls for this group demonstrate lower CBLM-cortical connectivity in frontal, temporal, parietal, and occipital cortices. Further, late postmenopausal male controls also exhibit lower CBLM-subcortical connectivity in areas including the caudate, putamen, amygdala, cingulate cortex, and insula, along with several areas of lower connectivity within the CBLM. On the other hand, late postmenopausal male controls show greater connectivity in some parietal areas, such as the precuneus and angular gyrus, as well as the corona radiata and thalamic radiation, also complementing some of the observed differences between relative female groups.

**Figure 4.**
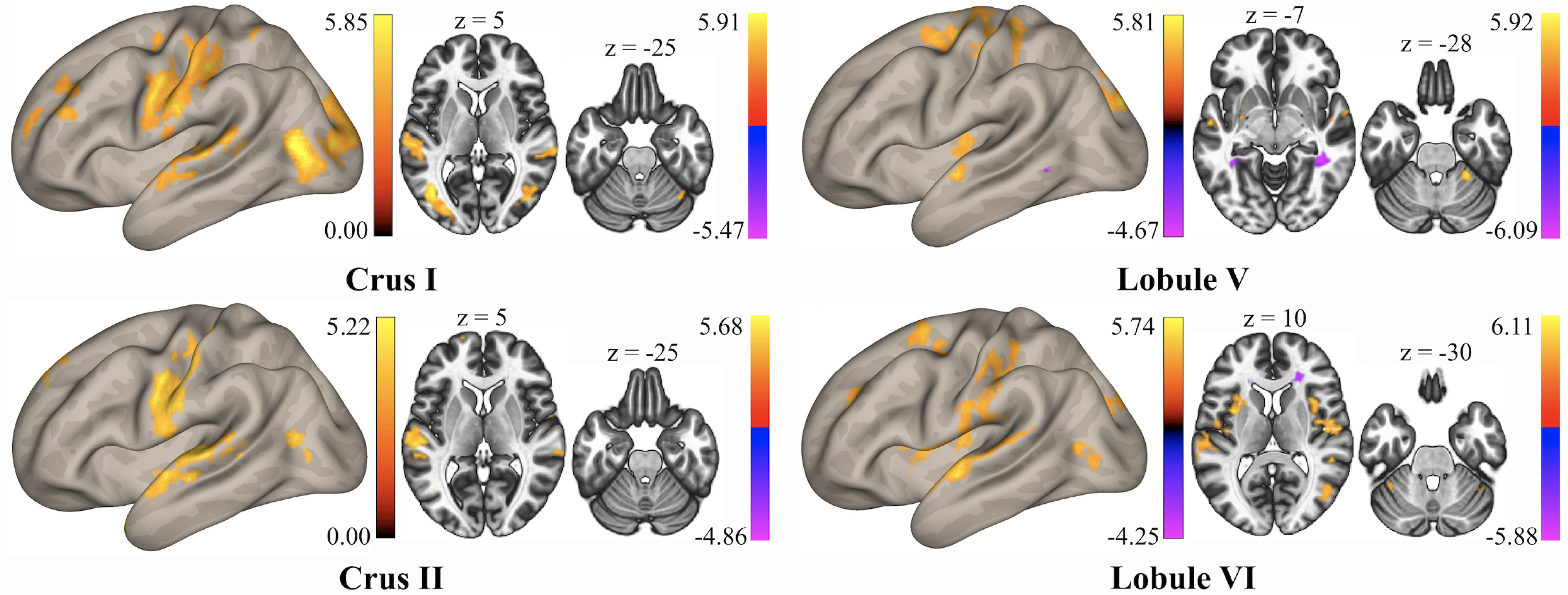
Reproductive male controls > late postmenopausal male controls. Contrasts between age-matched male controls for reproductive and late postmenopausal groups for Crus I & II (left) and Lobule V & VI (right). Orange/ yellow indicates greater connectivity in reproductive male controls, and blue/purple indicates greater connectivity in late postmenopausal male controls.

Nonetheless, in comparison to reproductive and late postmenopausal female groups, a few distinct differences emerge between male control groups. For instance, late postmenopausal male controls demonstrate lower connectivity in the thalamus, along with greater connectivity in the amygdala, relative to reproductive male controls. Likewise, late postmenopausal females show greater connectivity between several CBLM seeds and middle occipital areas, whereas the relative males only demonstrate lower connectivity with the occipital cortex. Notably, our comparisons between females and males are qualitative in nature; thus, results concerning sexspecific differences in CBLM network connectivity must be interpreted with caution. However, for a more direct comparison, we included a *post hoc* analysis investigating interactions between reproductive females, late postmenopausal females, reproductive male controls, and late postmenopausal male controls. These results can be found in **Supplementary Table 6**. In brief, this analysis reveals greater connectivity differences between reproductive and late postmenopausal females for Crus I/Crus II and middle temporal areas, whereas male control groups exhibit greater differences between Crus I/Crus II and the somatosensory cortex (**Figure 5**). Significant differences were not found for Lobule V or VI with this particular comparison.

**Figure 5.**
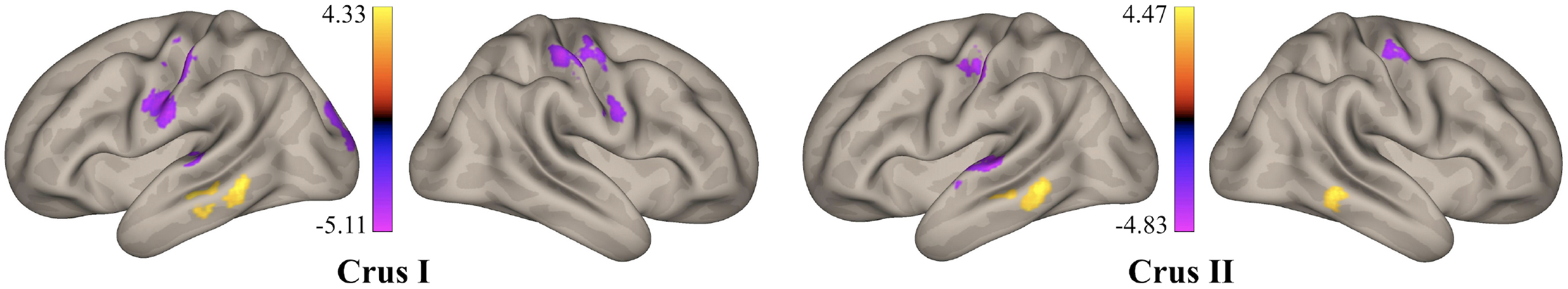
Female reproductive/late postmenopausal differences > relative male control differences. Interactions between reproductive and late postmenopausal females and relative age-matched male control groups for Crus I (left) and Crus II (right). Orange/yellow indicates greater connectivity differences between female groups, and blue/purple indicates greater differences between male groups.

## 4. Discussion

### 4.1. Summary

When investigating differences in CBLM resting state network connectivity between female reproductive stages, we found that late postmenopausal females primarily show lower CBLM-frontal and CBLM-striatal connectivity, as well as lower connectivity within the CBLM, compared to reproductive females. However, late postmenopausal females also demonstrate greater connectivity between the CBLM and a few cortical and subcortical areas. Some of these differences begin to emerge in transitional stages, such as perimenopause and early postmenopause. Further, relative comparisons between age-matched male controls display distinct connectivity patterns, though some similarities between sexes were discovered as well.

### 4.2. Impacts of Menopause on Cerebellar Networks

Given the observed differences in CBLM connectivity between reproductive and postmenopausal females, we suggest that menopause and the associated decreases in estrogen influence CBLM network differences in aging females. These reproductive stage differences broadly parallel general age differences in CBLM connectivity, as reported in the literature. Both cortical and subcortical connectivity with the CBLM is disrupted in older adults relative to younger (Bernard et al., 2013; Hausman et al., 2020), and we show similar patterns in postmenopausal females compared to reproductive. However, **Figure 1** displays an overlap in age range across female reproductive stages, even between reproductive and late postmenopausal groups. This suggests that age varies between female groups, which helps to offset inherent age impacts. Therefore, we propose that the current findings are predominantly a product of menopause and the accompanied hormonal changes, rather than solely age.

Taking into account the relationship between menopause, estrogen, and behavior (Daniel, 2013; Duka et al., 2000; Fischer et al., 2014; Gurvich et al., 2018; LeBlanc et al., 2001; Weber et al., 2014, 2013, 2012), changes in reproductive stage over the course of the female lifespan may contribute to elevated functional declines in aging females. Relative to males, females experience more severe cognitive and motor deficits with age (Hartholt et al., 2011; Kim et al., 2010; Panzer et al., 1995; Proust-Lima et al., 2008; Stevens and Sogolow, 2005), as well as higher incidence and severity of AD (Carter et al., 2012; Hebert et al., 2013; Mazure and Swendsen, 2016). To this point, the CBLM is implicated in AD (Tabatabaei-Jafari et al., 2017) and has a large role in behaviors disrupted with age, including balance, timing and execution of movement, processing speed, and working memory (Bernard and Seidler, 2014, 2013, Chen and Desmond, 2005a, 2005b; Eckert et al., 2010; Ivry et al., 2002, 1988; Morton and Bastian, 2004; Sullivan et al., 2010). In addition, the frontal cortex is heavily involved in memory and cognition (McAndrews and Milner, 1991; Paul et al., 2009), and the striatum regulates visuomotor skill, sequential patterns, and automaticity (Aizenstein et al., 2004; Doyon et al., 1997; Ungerleider et al., 2002). In the current work, we observed lower CBLM-frontal and CBLM-striatal connectivity across seeds in postmenopausal females, relative to both reproductive and perimenopausal females, and these patterns were different in males. Thus, altered CBLM network connectivity with menopause, as observed here, may exacerbate functional declines in aging females and elucidate potential biological mechanisms underlying sex differences in aging declines.

Pertinent to this idea, several works highlight sex-specific differences in the aging brain and CBLM volume, in particular (Raz et al., 2001, 1998; Rhyu et al., 1999; Ruigrok et al., 2014), though little work investigates CBLM network connectivity. We began to address this gap in prior work unveiling sex differences in ventral and dorsal dentate networks, as well as distinct associations with age in these CBLM networks between females and males (Bernard et al., 2021). The current investigation builds upon our past work, while continuing to fill the gap, by evaluating connectivity differences between females and males across multiple CBLM lobules.

Further, the current study examines CBLM connectivity through both cognitive and motor domains, providing additional context for potential applications to behavior. Considering evidence that estrogen fluctuations alter resting state connectivity (Petersen et al., 2014; Pletzer et al., 2016), it is likely that menopause contributes to more pronounced differences in CBLM connectivity in older females, compared to males. Thus, sex differences in the aging brain and, in turn, functional consequences with age may be partially attributed to this biological process. However, more work is needed to fully grasp the impact of menopause on the trajectory of brain changes within aging females.

Though unanticipated, our results demonstrate that postmenopausal females also exhibit greater CBLM connectivity than reproductive females with a few alternative areas. It is possible that these findings are brought about by compensatory scaffolding in the aging brain, which has been proposed as an adaptive response to age-related declines (Cabeza et al., 2018; Park and Reuter-Lorenz, 2009; Reuter-Lorenz and Park, 2014). Notably, however, greater connectivity is also observed in postmenopausal male controls, indicating that such differences may be common outcomes in aging more broadly. That is, scaffolding occurs in both sexes. Nonetheless, our explanations are entirely based on conjecture as our study was not tailored towards investigating these specific theories. Further, though we anticipated that connectivity differences would begin to emerge in the perimenopausal stage, we did not find any significant differences between reproductive and perimenopausal females. This could be explained to some degree by hormonal variability in the perimenopausal stage, as changes in sex hormones begin to occur. In fact, works by Jacobs et al. (2017) and Pritschet et al. (2020) highlight associations between hormonal fluctuations and functional network organization. However, without access to direct hormone data for the current sample, we cannot appropriately account for hormonal variability during the menopausal transition in our analyses.

Finally, our *post hoc* analysis, which directly compared differences between reproductive and late postmenopausal females to differences between age-matched male controls, yielded interesting results. First, female groups demonstrated greater differences in connectivity than their male counterparts between Crus I/Crus II and the middle temporal lobe. Atrophy of middle temporal areas, such as the hippocampus, is heavily associated with early AD (Busatto et al., 2003; Convit et al., 2000).(Busatto et al., 2003; Convit et al., 2000; Leal and Yassa, 2013). In fact, temporal stimulation improves visual recognition memory in AD patients (Boggio et al., 2012, 2009). Thus, given the increased incidence and symptomology of AD in females (Carter et al., 2012; Mazure and Swendsen, 2016), this finding could be particularly important for understanding sex differences in AD and other dementias. Second, compared to females, relative male controls demonstrate greater connectivity differences between Crus I/Crus II and the somatosensory cortex. This region is involved in the integration of sensory and motor information (Borich et al., 2015), which is essential for skilled movement, and could be especially pertinent to age-related motor declines and differences.

### 4.3. Limitations

Importantly, we did not have access to behavioral data. Functional relevance of the current findings is purely speculative. Moreover, we did not have access to direct hormone data, so it is possible that subjects taking hormonal contraceptives or undergoing hormone therapy could have been included in our sample. Relatedly, classification of female reproductive stage was based only on self-reported information, due to the lack of hormone data. In addition, males and females were not directly compared in our primary analyses, though we did complete a *post hoc* analysis of interactions (see **Supplementary Table 6**). As such, many of our sex comparisons are qualitative in nature. We chose to adopt this statistical approach as our principal objective was to evaluate the influence of reproductive stage on CBLM networks within aging females, in particular. The fundamental purpose of our relative male comparisons was to establish an indirect reference for results from the female comparisons. Finally, as menopause is inherently tied to age, we cannot completely discount the impact of age on our results. However, we also cannot adequately control for age in our analyses as such an approach would incidentally factor out the effect of menopause. To reiterate, we have illustrated significant age overlap between female groups (**Figure 1**), indicating effective categorization of females based on reproductive stage in lieu of simply age. Comparisons between age-matched males further rule out impacts solely due to age and emphasize the effect of reproductive stage on the female brain.

### 4.4. Conclusions

The current study enhances our understanding of the relationship between female reproductive stage, and accompanied differences in estrogen, and resting state CBLM network connectivity across adulthood. There is limited aging research targeting CBLM connectivity in the context of menopause, a key component of aging in females; however, our results indicate that this relatively understudied factor modulates connectivity between the CBLM and both cortical and subcortical areas. We suggest that menopause contributes to differences in age-related declines within females, as well as between sexes, and should be investigated further in future aging research. Moreover, little work adopts a regional or lobular approach when investigating CBLM network connectivity. However, our findings show regional differences between cognitive- and motor-focused CBLM lobules and other areas of the brain, illustrating the importance of a more localized perspective on CBLM connectivity in aging research. In light of the individual and societal consequences of aging, it is important to probe the mechanisms underlying brain differences in advanced age, and their unique course in females and males, to better combat functional declines and diseases, such as AD, in older adulthood.

## Acknowledgements

Data for this investigation was acquired from the Cambridge Centre for Ageing and Neuroscience (Cam-CAM) repository. Funding for the Cam-CAN study was provided by the UK Biotechnology and Biological Sciences Research Council (grant number BB/H008217/1); the University of Cambridge, UK; and the UK Medical Research Council. The current work was further supported by the National Institute on Aging (grant number R01AG065010).

## Notes

https://www.cam-can.org/index.php?content=dataset

### Competing Interest Statement

The authors have declared no competing interest.

