## Supplemental Tables for "The influence of reproductive stage on cerebellar network connectivity across adulthood"

| **Region** | **Cluster Size** | **MNI Coordinates** | | | **P_(FDR)_** | **Beta** |
| --- | --- | --- | --- | --- | --- | --- |
|  |  | **X** | **Y** | **Z** |  |  |
| **REPRODUCTIVE FEMALES > LATE POSTMENOPAUSAL FEMALES** | | | | | | |
| **Right Crus I** | | | | | | |
| Anterior Limb of Internal Capsule, Left | 1143 | -12 | 8 | 8 | 0.000 | 0.096 |
| Middle Frontal Gyrus (Posterior Segment), Left | 846 | -48 | 14 | 46 | 0.000 | 0.120 |
| Crus I of Cerebellum, Right | 831 | 36 | -74 | -34 | 0.000 | 0.150 |
| Supramarginal Gyrus, Left | 796 | -62 | -48 | 42 | 0.000 | 0.130 |
| Anterior Limb of Internal Capsule, Right | 476 | 14 | 2 | 16 | 0.000 | 0.096 |
| Lateral Ventricle (Frontal), Right | 398 | 14 | 26 | -4 | 0.000 | -0.097 |
| Superior Corona Radiata, Left | 360 | -24 | -16 | 26 | 0.000 | -0.078 |
| Anterior Corona Radiata, Left | 223 | -16 | 32 | -4 | 0.000 | -0.088 |
| Superior Frontal Gyrus (Prefrontal Cortex), Left | 222 | -10 | 46 | 40 | 0.000 | 0.110 |
| Inferior Frontal Gyrus (Pars Opercularis), Left | 150 | -40 | 16 | 24 | 0.001 | 0.092 |
| Cerebellar Lobule VI, Left | 143 | -22 | -72 | -22 | 0.001 | 0.130 |
| Superior Frontal Gyrus (Prefrontal Cortex), Left | 138 | -20 | 62 | 26 | 0.001 | 0.120 |
| Splenium of Corpus Callosum, Right | 117 | 14 | -34 | 12 | 0.002 | -0.091 |
| Superior Frontal Gyrus (Posterior Segment), Left | 86 | -12 | 32 | 54 | 0.010 | 0.120 |
| Superior Parietal Gyrus, Left | 85 | -28 | -54 | 68 | 0.010 | 0.110 |
| Superior Corona Radiata, Right | 76 | 28 | 0 | 26 | 0.015 | -0.078 |
| Inferior Frontal Gyrus (Pars Orbitralis),  Left | 73 | -42 | 34 | -14 | 0.017 | 0.100 |
| Middle Occipital Gyrus, Right | 71 | 30 | -68 | 8 | 0.017 | -0.078 |
| Crus I of Cerebellum, Right | 71 | 6 | -80 | -22 | 0.017 | 0.160 |
| Posterior Thalamic Radiation, Left | 68 | -32 | -54 | 2 | 0.018 | -0.084 |
| Amygdala, Right | 68 | 28 | -6 | -16 | 0.018 | 0.079 |
| Inferior Temporal Gyrus, Left | 66 | -46 | 4 | -36 | 0.019 | 0.093 |
| **Right Crus II** | | | | | | |
| Inferior Frontal Gyrus (Pars Triangularis),  Left | 1544 | -34 | 20 | 20 | 0.000 | 0.120 |
| Lateral Occipital Cortex (Superior Division), Left | 861 | -44 | -70 | 50 | 0.000 | 0.140 |
| Caudate Nucleus, Left | 831 | -12 | 2 | 14 | 0.000 | 0.097 |
| Crus I of Cerebellum, Right | 803 | 6 | -80 | -24 | 0.000 | 0.170 |
| Middle Temporal Gyrus, Left | 514 | -68 | -44 | -14 | 0.000 | 0.120 |
| Inferior Frontal Gyrus (Pars Orbitralis),  Left | 376 | -46 | 44 | -8 | 0.000 | 0.120 |
| Superior Longitudinal Fasciculus, Left | 375 | -32 | -28 | 28 | 0.000 | -0.077 |
| Middle Occipital Gyrus, Left | 370 | -28 | -94 | -12 | 0.000 | -0.091 |
| Lateral Ventricle (Frontal), Right | 348 | 10 | 4 | 14 | 0.000 | 0.094 |
| Superior Parietal Gyrus, Right | 197 | 20 | -38 | 38 | 0.000 | -0.094 |
| Caudate Nucleus, Right | 163 | 12 | 24 | -4 | 0.000 | -0.097 |
| Superior Parietal Gyrus, Right | 131 | 18 | -64 | 38 | 0.001 | -0.086 |
| Superior Frontal Gyrus (Prefrontal  Cortex), Left | 122 | -12 | 44 | 38 | 0.002 | 0.120 |
| Posterior Thalamic Radiation, Right | 98 | 36 | -58 | 4 | 0.005 | -0.086 |
| Inferior Frontal Gyrus (Pars  Triangularis), Right | 73 | 42 | 28 | 12 | 0.020 | 0.098 |
| Dorsal Anterior Cingulate Gyrus, Left | 67 | -6 | -4 | 38 | 0.025 | -0.092 |
| Posterior Thalamic Radiation, Left | 67 | -32 | -62 | 2 | 0.025 | -0.084 |
| Crus I of Cerebellum, Left | 66 | -24 | -80 | -32 | 0.025 | 0.110 |
| Cerebellar Lobule VIII, Right | 65 | 24 | -44 | -54 | 0.025 | -0.100 |
| Precuneus Cortex | 59 | -2 | -82 | 42 | 0.034 | -0.110 |
| **Right Lobule V** | | | | | | |
| Amygdala, Left | 1979 | -24 | -6 | -14 | 0.000 | 0.090 |
| Cerebellar Lobule IV-V, Right | 1571 | 20 | -42 | -24 | 0.000 | 0.110 |
| Precentral Gyrus, Left | 830 | -38 | -20 | 50 | 0.000 | 0.098 |
| Dorsal Anterior Cingulate Gyrus, Left | 826 | -4 | -4 | 42 | 0.000 | 0.093 |
| Corticospinal Tract, Right | 629 | 16 | -12 | -16 | 0.000 | 0.096 |
| Superior Frontal Gyrus (Dorsolateral),  Right | 216 | 20 | -16 | 74 | 0.000 | 0.092 |
| Supramarginal Gyrus, Right | 213 | 42 | -24 | 20 | 0.000 | 0.089 |
| Middle Occipital Gyrus, Left | 163 | -32 | -60 | 34 | 0.000 | -0.079 |
| Putamen, Right | 156 | 26 | 8 | 6 | 0.000 | 0.082 |
| Postcentral Gyrus, Left | 139 | -58 | -4 | 46 | 0.001 | 0.087 |
| External Capsule, Right | 127 | 30 | -14 | 8 | 0.001 | 0.081 |
| Middle Frontal Gyrus (Dorsal Prefrontal Cortex), Left | 125 | -28 | 54 | -8 | 0.001 | -0.090 |
| Posterior Insula, Right | 103 | 40 | -4 | 4 | 0.003 | 0.085 |
| Anterior Corona Radiata, Left | 97 | -16 | 28 | -6 | 0.003 | -0.082 |
| Anterior Corona Radiata, Right | 81 | 22 | 36 | -4 | 0.008 | -0.077 |
| Angular Gyrus, Right | 80 | 52 | -44 | 36 | 0.008 | -0.078 |
| Posterior Middle Temporal Gyrus, Left | 68 | -44 | -40 | -6 | 0.015 | -0.082 |
| Postcentral Gyrus, Right | 66 | 38 | -16 | 40 | 0.016 | 0.082 |
| Superior Corona Radiata, Right | 58 | 24 | 2 | 30 | 0.023 | -0.070 |
| Pole of Superior Temporal Gyrus, Right | 58 | 56 | 2 | -8 | 0.023 | 0.089 |
| Postcentral Gyrus, Right | 53 | 62 | 2 | 34 | 0.031 | 0.084 |
| Lateral Ventricle, Right | 50 | 6 | 2 | 20 | 0.036 | 0.085 |
| **Right Lobule VI** | | | | | | |
| Cerebellar Lobule VI, Right | 1887 | 20 | -72 | -18 | 0.000 | 0.130 |
| Amygdala, Left | 865 | -24 | -6 | -14 | 0.000 | 0.096 |
| Supramarginal Gyrus, Left | 794 | -56 | -24 | 28 | 0.000 | 0.097 |
| Middle Occipital Gyrus, Left | 602 | -44 | -76 | 20 | 0.000 | 0.099 |
| Amygdala, Right | 549 | 26 | -8 | -14 | 0.000 | 0.096 |
| Middle Occipital Gyrus, Right | 506 | 40 | -80 | 18 | 0.000 | 0.100 |
| Putamen, Left | 453 | -22 | 12 | -2 | 0.000 | 0.092 |
| Putamen, Right | 435 | 24 | 6 | 6 | 0.000 | 0.084 |
| Precentral Gyrus, Left | 356 | -60 | 4 | 30 | 0.000 | 0.100 |
| Angular Gyrus, Right | 343 | 50 | -46 | 48 | 0.000 | -0.094 |
| Postcentral Gyrus, Right | 280 | 62 | -18 | 42 | 0.000 | 0.097 |
| Precentral Gyrus, Left | 272 | -28 | -6 | 52 | 0.000 | 0.097 |
| Cerebellar Dentate Nucleus, Left | 268 | -16 | -60 | -34 | 0.000 | 0.099 |
| Precentral Gyrus, Right | 260 | 62 | 6 | 32 | 0.000 | 0.110 |
| Anterior Corona Radiata, Right | 186 | 22 | 36 | -2 | 0.000 | -0.085 |
| Anterior Corona Radiata, Left | 150 | -16 | 34 | -2 | 0.001 | -0.092 |
| Sagittal Stratum, Left | 132 | -44 | -38 | -8 | 0.001 | -0.087 |
| Superior Parietal Gyrus, Left | 132 | -26 | -52 | 70 | 0.001 | 0.110 |
| Middle Frontal Gyrus, Right | 129 | 38 | 62 | -6 | 0.001 | -0.100 |
| Superior Parietal Gyrus, Right | 118 | 24 | -58 | 70 | 0.002 | 0.098 |
| Sagittal Stratum, Right | 113 | 42 | -38 | -6 | 0.002 | -0.087 |
| Dorsal Anterior Cingulate Gyrus, Left | 101 | -6 | 0 | 46 | 0.004 | 0.094 |
| Precentral Gyrus, Right | 68 | 36 | -2 | 26 | 0.021 | -0.071 |
| Lingual Gyrus, Right | 67 | 28 | -56 | -4 | 0.021 | 0.098 |
| Superior Corona Radiata, Left | 58 | -24 | -10 | 26 | 0.033 | -0.072 |
| Middle Frontal Gyrus (Dorsal Prefrontal Cortex), Left | 58 | -28 | 54 | -6 | 0.033 | -0.093 |
| Superior Temporal Gyrus, Left | 55 | -50 | 0 | 0 | 0.037 | 0.098 |

**Supplementary Table 1.** *Cerebellar network connectivity between reproductive and late postmenopausal females across seeds.* Anatomical regions were identified using the Johns Hopkins University Atlas and the Automated Anatomical Labeling Atlas, Version 3 (Rolls et al., 2020), in MRIcron. Cerebellar deep nuclei were identified using the atlas provided by Dimitrova et al. (2002). Beta values are unstandardized.

| **Region** | **Cluster Size** | **MNI Coordinates** | | | **P_(FDR)_** | **Beta** |
| --- | --- | --- | --- | --- | --- | --- |
|  |  | **X** | **Y** | **Z** |  |  |
| **REPRODUCTIVE FEMALES > EARLY POSTMENOPAUSAL FEMALES** | | | | | | |
| **Right Crus I** | | | | | | |
| Precuneus, Right | 127 | 2 | -44 | 50 | 0.003 | 0.140 |
| Precentral Gyrus, Left | 125 | -30 | -6 | 54 | 0.003 | 0.140 |
| Superior Fronto-Occipital Fasciculus,  Right | 95 | 22 | 10 | 20 | 0.011 | -0.120 |
| **Right Crus II** | | | | | | |
| Fusiform Gyrus, Left | 182 | -16 | -88 | -16 | 0.000 | -0.140 |
| Inferior Occipital Gyrus, Right | 95 | 38 | -96 | -6 | 0.011 | -0.150 |
| Cuneus, Left | 87 | -12 | -96 | 26 | 0.012 | -0.140 |
| **Right Lobule V** | | | | | | |
| Amygdala, Right | 197 | 24 | -10 | -12 | 0.000 | 0.130 |
| Putamen, Left | 176 | -26 | 0 | 2 | 0.000 | 0.130 |
| Parahippocampal Gyrus, Left | 119 | -12 | -10 | -18 | 0.002 | 0.130 |
| Putamen, Right | 62 | 28 | -12 | 6 | 0.034 | 0.130 |
| Parahippocampal Gyrus, Right | 62 | 20 | -26 | -10 | 0.034 | 0.130 |
| **Right Lobule VI** | | | | | | |
| Amygdala, Left | 221 | -20 | -10 | -18 | 0.000 | 0.150 |
| Hippocampus, Right | 172 | 26 | -10 | -16 | 0.000 | 0.130 |

**Supplementary Table 2.** *Cerebellar network connectivity between reproductive and early postmenopausal females across seeds.* Anatomical regions were identified using the Johns Hopkins University Atlas and the Automated Anatomical Labeling Atlas, Version 3 (Rolls et al., 2020), in MRIcron. Beta values are unstandardized.

| **Region** | **Cluster Size** | **MNI Coordinates** | | | **P_(FDR)_** | **Beta** |
| --- | --- | --- | --- | --- | --- | --- |
|  |  | **X** | **Y** | **Z** |  |  |
| **PERIMENOPAUSAL FEMALES > LATE POSTMENOPAUSAL FEMALES** | | | | | | |
| **Right Crus I** | | | | | | |
| Caudate, Left | 776 | -12 | 8 | 10 | 0.000 | 0.130 |
| Anterior Limb of Internal Capsule, Right | 477 | 14 | 10 | 2 | 0.000 | 0.130 |
| **Right Crus II** | | | | | | |
| Inferior Frontal Gyrus (Pars Opercularis), Left | 437 | -54 | 16 | 22 | 0.000 | 0.160 |
| Inferior Frontal Gyrus (Pars Orbitralis), Left | 168 | -42 | 40 | 2 | 0.001 | 0.160 |
| Putamen, Left | 165 | -26 | -6 | -2 | 0.001 | 0.110 |
| Caudate, Left | 148 | -12 | 10 | 8 | 0.002 | 0.140 |
| Caudate, Right | 145 | 12 | 10 | 6 | 0.002 | 0.120 |
| **Right Lobule V** | | | | | | |
| Superior Temporal Gyrus, Left | 123 | -46 | -12 | 6 | 0.009 | 0.120 |
| Cerebellar Lobule VI, Right | 95 | 32 | -34 | -34 | 0.018 | 0.170 |
| Dorsal Anterior Cingulate Gyrus, Right | 81 | 8 | -8 | 40 | 0.018 | 0.130 |
| Posterior Cingulate Gyrus, Right | 80 | 0 | -48 | 10 | 0.018 | 0.140 |
| Postcentral Gyrus, Left | 75 | -42 | -20 | 52 | 0.018 | 0.140 |
| Postcentral Gyrus, Left | 73 | -42 | -10 | 18 | 0.018 | 0.120 |
| White Matter, Brainstem | 72 | -12 | -12 | -18 | 0.018 | 0.120 |
| **Right Lobule VI** | | | | | | |
| Putamen, Left | 133 | -18 | 10 | -2 | 0.013 | 0.120 |

**Supplementary Tale 3.** *Cerebellar network connectivity between perimenopausal and late postmenopausal females across seeds.* Anatomical regions were identified using the Johns Hopkins University Atlas and the Automated Anatomical Labeling Atlas, Version 3 (Rolls et al., 2020), in MRIcron. Beta values are unstandardized.

| **Region** | **Cluster Size** | **MNI Coordinates** | | | **P_(FDR)_** | **Beta** |
| --- | --- | --- | --- | --- | --- | --- |
|  |  | **X** | **Y** | **Z** |  |  |
| **EARLY POSTMENOPAUSAL FEMALES > LATE POSTMENOPAUSAL FEMALES** | | | | | | |
| **Right Crus I** | | | | | | |
| Superior Frontal Gyrus (Prefrontal Cortex), Left | 161 | -6 | 38 | 54 | 0.001 | 0.180 |
| Crus II of Cerebellum, Right | 156 | 30 | -80 | -36 | 0.001 | 0.240 |
| Postcentral Gyrus, Left | 150 | -16 | -36 | 48 | 0.001 | -0.130 |
| Superior Frontal Gyrus (Posterior  Segment), Right | 137 | 8 | -6 | 64 | 0.001 | -0.120 |
| Precuneus, Right | 121 | 2 | -44 | 50 | 0.001 | -0.130 |
| Posterior Cingulate Gyrus, Right | 121 | 12 | -36 | 38 | 0.001 | -0.130 |
| Middle Temporal Gyrus, Left | 81 | -62 | -22 | -12 | 0.011 | 0.150 |
| Superior Temporal Gyrus, Right | 56 | 64 | -24 | 14 | 0.046 | -0.130 |
| **Right Crus II** | | | | | | |
| Superior Parietal Gyrus, Right | 157 | 18 | -36 | 44 | 0.001 | -0.130 |
| Angular Gyrus, Left | 138 | -54 | -48 | 24 | 0.001 | 0.170 |
| Inferior Frontal Gyrus (Pars Orbitralis),  Left | 137 | -46 | 32 | -8 | 0.001 | 0.160 |
| Middle Temporal Gyrus, Left | 126 | -70 | -36 | -6 | 0.002 | 0.160 |
| Dorsal Anterior Cingulate Gyrus, Left | 114 | -6 | 0 | 40 | 0.003 | -0.130 |
| Superior Occipital Gyrus, Right | 89 | 18 | -60 | 30 | 0.008 | -0.120 |
| Superior Frontal Gyrus (Frontal Pole),  Right | 89 | 0 | 66 | 2 | 0.008 | 0.160 |
| Postcentral Gyrus, Left | 88 | -12 | -38 | 54 | 0.008 | -0.120 |
| Lateral Ventricle (Frontal), Right | 83 | 8 | 6 | 4 | 0.009 | 0.130 |
| Superior Frontal Gyrus (Medial), Left | 74 | -6 | 48 | 50 | 0.014 | 0.180 |
| Superior Frontal Gyrus (Posterior  Segment), Right | 57 | 4 | -6 | 66 | 0.037 | -0.120 |
| Cerebellar Lobule VI, Right | 54 | 30 | -76 | -24 | 0.042 | 0.210 |
| **Right Lobule V** | | | | | | |
| Middle Frontal Gyrus (Orbital), Right | 131 | 42 | 60 | -8 | 0.003 | -0.130 |
| Superior Frontal Gyrus (Frontal Pole),  Left | 96 | -10 | 64 | 2 | 0.011 | -0.110 |
| **Right Lobule VI** | | | | | | |
| Cerebellar Lobule IV-V, Right | 91 | 18 | -48 | -14 | 0.019 | 0.180 |
| Middle Frontal Gyrus (Dorsal Prefrontal  Cortex), Right | 89 | 38 | 60 | -8 | 0.019 | -0.130 |

**Supplementary Table 4.** *Cerebellar network connectivity between early and late postmenopausal females across seeds.* Anatomical regions were identified using the Johns Hopkins University Atlas and the Automated Anatomical Labeling Atlas, Version 3 (Rolls et al., 2020), in MRIcron. Beta values are unstandardized.

| **Region** | **Cluster Size** | **MNI Coordinates** | | | **P_(FDR)_** | **Beta** |
| --- | --- | --- | --- | --- | --- | --- |
|  |  | **X** | **Y** | **Z** |  |  |
| **REPRODUCTIVE MALE CONTROLS > LATE POSTMENOPAUSAL  MALE CONTROLS** | | | | | | |
| **Right Crus I** | | | | | | |
| Postcentral Gyrus, Left | 1802 | -60 | -12 | 24 | 0.000 | 0.110 |
| Postcentral Gyrus, Right | 1438 | 56 | -6 | 26 | 0.000 | 0.110 |
| Middle Occipital Gyrus, Left | 1223 | -42 | -72 | 6 | 0.000 | 0.110 |
| Superior Temporal Gyrus, Left | 836 | -52 | -18 | -4 | 0.000 | 0.110 |
| Middle Occipital Gyrus, Right | 734 | 30 | -76 | 24 | 0.000 | 0.110 |
| Inferior Occipital Gyrus, Right | 483 | 44 | -68 | -2 | 0.000 | 0.110 |
| Superior Temporal Gyrus, Right | 315 | 68 | -34 | 8 | 0.000 | 0.100 |
| Middle Frontal Gyrus (Dorsal Prefrontal  Cortex), Left | 265 | -26 | 38 | 40 | 0.000 | 0.110 |
| Superior Temporal Gyrus, Right | 265 | 44 | -4 | -18 | 0.000 | 0.100 |
| Cerebellar Lobule IV-V, Left | 178 | -16 | -48 | -12 | 0.000 | 0.093 |
| Lingual Gyrus, Right | 152 | 18 | -74 | -6 | 0.000 | 0.098 |
| Putamen, Left | 94 | -22 | 8 | -6 | 0.005 | 0.081 |
| Middle Frontal Gyrus (Posterior  Segment), Right | 91 | 36 | 32 | 36 | 0.006 | 0.100 |
| Lingual Gyrus, Right | 88 | 22 | -52 | -8 | 0.006 | 0.088 |
| Caudate Nucleus, Left | 85 | -10 | 10 | 12 | 0.007 | 0.110 |
| Superior Corona Radiata, Right | 81 | 24 | -4 | 26 | 0.008 | -0.089 |
| Superior Parietal Gyrus, Left | 74 | -34 | -48 | 68 | 0.011 | 0.120 |
| Middle Frontal Gyrus (Dorsal Prefrontal  Cortex), Left | 74 | -28 | 56 | 16 | 0.011 | 0.110 |
| Cerebellar Lobule VI, Right | 72 | 40 | -52 | -26 | 0.011 | 0.150 |
| Superior Corona Radiata, Right | 71 | 28 | -18 | 32 | 0.011 | -0.078 |
| Amygdala, Right | 71 | 24 | -10 | -16 | 0.011 | 0.092 |
| Superior Occipital Gyrus, Right | 68 | 16 | -68 | 22 | 0.013 | 0.099 |
| Nucleus Accumbens, Left | 59 | -12 | 16 | -10 | 0.022 | 0.082 |
| Middle Frontal Gyrus (Dorsal Prefrontal  Cortex), Left | 58 | -42 | 40 | 22 | 0.023 | 0.100 |
| Superior Corona Radiata, Left | 51 | -26 | -10 | 32 | 0.034 | -0.071 |
| Spinal Cord | 50 | -8 | -20 | -52 | 0.035 | -0.086 |
| Superior Frontal Gyrus (Posterior  Segment), Left | 46 | -22 | -6 | 52 | 0.045 | 0.093 |
| **Right Crus II** | | | | | | |
| Superior Temporal Gyrus, Left | 1765 | -58 | -24 | 4 | 0.000 | 0.110 |
| Superior Frontal Gyrus (Posterior  Segment), Right | 400 | 14 | 36 | 52 | 0.000 | 0.120 |
| Postcentral Gyrus, Right | 393 | 58 | -4 | 26 | 0.000 | 0.110 |
| Caudate Nucleus, Left | 246 | -10 | 10 | 12 | 0.000 | 0.100 |
| Superior Temporal Gyrus, Right | 199 | 64 | -20 | 0 | 0.000 | 0.100 |
| Superior Corona Radiata, Left | 139 | -26 | -10 | 32 | 0.001 | -0.074 |
| Precentral Gyrus, Right | 134 | 38 | -18 | 50 | 0.001 | 0.097 |
| Superior Frontal Gyrus (Frontal Pole),  Left | 103 | -20 | 68 | 6 | 0.006 | 0.120 |
| Pole of Superior Temporal Gyrus, Left | 82 | -40 | 18 | -30 | 0.014 | 0.100 |
| Cerebellar Lobule VIII, Right | 82 | 28 | -66 | -42 | 0.014 | 0.150 |
| Amygdala, Left | 80 | -26 | -6 | -10 | 0.014 | 0.082 |
| Amygdala, Right | 79 | 26 | -2 | -12 | 0.014 | 0.089 |
| Precentral Gyrus, Left | 76 | -4 | -26 | 62 | 0.016 | 0.096 |
| Caudate Nucleus, Right | 74 | 12 | 6 | 18 | 0.016 | 0.110 |
| Spinal Cord | 67 | -10 | -22 | -54 | 0.023 | -0.092 |
| Precentral Gyrus, Right | 56 | 52 | -6 | 10 | 0.041 | 0.100 |
| Middle Occipital Gyrus, Left | 56 | -46 | -64 | 8 | 0.041 | 0.099 |
| **Right Lobule V** | | | | | | |
| Superior Parietal Gyrus, Left | 1413 | -28 | -36 | 58 | 0.000 | 0.099 |
| Dorsal Anterior Cingulate Gyrus, Right | 1211 | 0 | 2 | 36 | 0.000 | 0.094 |
| Superior Occipital Gyrus, Left | 487 | -16 | -88 | 26 | 0.000 | 0.095 |
| Postcentral Gyrus, Right | 407 | 40 | -10 | 20 | 0.000 | 0.089 |
| Inferior Fronto-Occipital Fasciculus, Left | 380 | -30 | -4 | -8 | 0.000 | 0.083 |
| Lobule IX of Cerebellar Vermis | 365 | 6 | -60 | -38 | 0.000 | 0.098 |
| Superior Occipital Gyrus, Right | 315 | 20 | -76 | 28 | 0.000 | 0.091 |
| Cerebellar Lobule IV-V, Right | 220 | 26 | -40 | -28 | 0.000 | 0.130 |
| Precentral Gyrus, Right | 211 | 32 | -8 | 56 | 0.000 | 0.090 |
| Anterior Corona Radiata, Right | 187 | 22 | 32 | 10 | 0.000 | -0.093 |
| Cingulum (Cingulate Gyrus), Right | 159 | 12 | -20 | 34 | 0.000 | 0.089 |
| Superior Temporal Gyrus, Left | 151 | -42 | -12 | 12 | 0.000 | 0.089 |
| Sagittal Stratum, Right | 149 | 40 | -40 | -6 | 0.000 | -0.094 |
| Posterior Thalamic Radiation, Left | 144 | -36 | -46 | -2 | 0.000 | -0.096 |
| Cerebral Peduncle, Left | 120 | -12 | -16 | -18 | 0.001 | 0.089 |
| Thalamus, Right | 113 | 12 | -6 | -16 | 0.001 | 0.090 |
| Anterior Limb of Internal Capsule, Right | 98 | 12 | 4 | 6 | 0.003 | 0.080 |
| Anterior Corona Radiata, Left | 95 | -20 | 22 | 12 | 0.003 | -0.088 |
| Superior Parietal Gyrus, Right | 95 | 34 | -36 | 52 | 0.003 | 0.090 |
| Superior Temporal Gyrus, Left | 75 | -56 | -10 | -8 | 0.008 | 0.091 |
| Supramarginal Gyrus, Right | 74 | 44 | -22 | 20 | 0.009 | 0.093 |
| Superior Fronto-Occipital Fasciculus,  Right | 68 | 20 | 4 | 24 | 0.012 | -0.081 |
| White Matter, Cerebellum | 65 | 24 | -36 | -50 | 0.014 | 0.088 |
| Cerebellar Lobule IV-V, Left | 53 | -22 | -34 | -32 | 0.029 | 0.100 |
| White Matter, Cerebellum | 50 | 8 | -48 | -62 | 0.034 | 0.079 |
| **Right Lobule VI** | | | | | | |
| Superior Temporal Gyrus, Left | 1667 | -58 | -10 | -6 | 0.000 | 0.017 |
| Superior Temporal Gyrus, Right | 1204 | 48 | -12 | 12 | 0.000 | 0.013 |
| Dorsal Anterior Cingulate Gyrus, Right | 1196 | 0 | 4 | 36 | 0.000 | -0.008 |
| Middle Occipital Gyrus, Right | 465 | 48 | -66 | 4 | 0.000 | 0.020 |
| Middle Occipital Gyrus, Left | 417 | -28 | -76 | 26 | 0.000 | 0.022 |
| Pole of Superior Temporal Gyrus, Right | 401 | 52 | 12 | -14 | 0.000 | -0.001 |
| Insular, Left | 300 | -36 | 4 | 10 | 0.000 | 0.049 |
| Cerebellar Lobule IV-V, Left | 260 | -4 | -62 | -6 | 0.000 | 0.150 |
| Anterior Corona Radiata, Right | 231 | 24 | 32 | 0 | 0.000 | -0.040 |
| Precuneus, Right | 194 | 8 | -72 | 32 | 0.000 | -0.024 |
| Precentral Gyrus, Right | 189 | 56 | 8 | 32 | 0.000 | 0.043 |
| Superior Frontal Gyrus (Posterior  Segment), Left | 166 | -18 | 6 | 62 | 0.000 | 0.008 |
| Dorsal Anterior Cingulate Gyrus, Right | 156 | 10 | -24 | 38 | 0.000 | -0.032 |
| Middle Frontal Gyrus (Dorsal Prefrontal  Cortex), Left | 143 | -30 | 42 | 24 | 0.001 | 0.037 |
| Cerebellar Lobule VI, Left | 120 | -36 | -52 | -26 | 0.002 | 0.290 |
| Middle Frontal Gyrus (Dorsal Prefrontal  Cortex), Right | 112 | 28 | 42 | 24 | 0.002 | 0.031 |
| Caudate Nucleus, Left | 111 | -14 | -6 | 20 | 0.002 | -0.007 |
| Cerebellar Lobule VI, Right | 108 | 24 | -54 | -20 | 0.002 | 0.480 |
| Superior Parietal Gyrus, Left | 101 | -16 | -30 | 38 | 0.003 | -0.033 |
| Middle Occipital Gyrus, Right | 99 | 26 | -74 | 38 | 0.003 | 0.037 |
| Crus I of Cerebellum, Right | 95 | 48 | -50 | -34 | 0.004 | 0.330 |
| Middle Occipital Gyrus, Right | 86 | 30 | -76 | 24 | 0.006 | 0.059 |
| Posterior Superior Temporal Gyrus,  Right | 75 | 52 | -40 | 12 | 0.011 | -0.020 |
| Genu of Corpus Callosum | 70 | -18 | 32 | 0 | 0.015 | -0.028 |
| Posterior Middle Temporal Gyrus, Right | 68 | 44 | -36 | -8 | 0.016 | -0.033 |
| Cerebellar Lobule VI, Left | 65 | -18 | -64 | -20 | 0.018 | 0.450 |
| Fusiform Gyrus, Right | 63 | 32 | -2 | -36 | 0.020 | -0.015 |
| Middle Occipital Gyrus, Left | 62 | -42 | -66 | 6 | 0.021 | 0.032 |
| Amygdala, Right | 57 | 26 | -4 | -16 | 0.027 | -0.002 |
| Angular Gyrus, Right | 52 | 44 | -58 | 50 | 0.036 | -0.044 |
| Inferior Fronto-Occipital Fasciculus, Left | 50 | -30 | -4 | -8 | 0.040 | 0.073 |

**Supplementary Table 5.** *Cerebellar network connectivity between age-matched reproductive male controls and late postmenopausal male controls across seeds*. Anatomical regions were identified using the Johns Hopkins University Atlas and the Automated Anatomical Labeling Atlas, Version 3 (Rolls et al., 2020), in MRIcron. Beta values are unstandardized.

| **Region** | **Cluster Size** | **MNI Coordinates** | | | **P_(FDR)_** | **Beta** |
| --- | --- | --- | --- | --- | --- | --- |
|  |  | **X** | **Y** | **Z** |  |  |
| **INTERACTION EFFECTS** | | | | | | |
| **Right Crus I** | | | | | | |
| Postcentral Gyrus, Right | 469 | 46 | -30 | 56 | 0.000 | -0.150 |
| Postcentral Gyrus, Left | 285 | -62 | -10 | 26 | 0.000 | -0.150 |
| Postcentral Gyrus, Left | 258 | -48 | -20 | 50 | 0.000 | -0.160 |
| Middle Occipital Gyrus, Left | 193 | -22 | -98 | 10 | 0.000 | -0.140 |
| Precuneus, Right | 161 | 14 | -66 | 24 | 0.000 | -0.140 |
| Middle Temporal Gyrus, Left | 103 | -60 | -24 | -12 | 0.003 | 0.150 |
| Postcentral Gyrus, Right | 101 | 54 | -4 | 26 | 0.003 | -0.140 |
| Middle Temporal Gyrus, Left | 89 | -62 | -46 | -10 | 0.006 | 0.150 |
| Superior Temporal Gyrus, Left | 63 | -48 | -24 | 2 | 0.026 | -0.140 |
| **Right Crus II** | | | | | | |
| Superior Temporal Gyrus, Left | 217 | -68 | -20 | 4 | 0.000 | -0.150 |
| Postcentral Gyrus, Left | 215 | -46 | -14 | 52 | 0.000 | -0.150 |
| Middle Temporal Gyrus, Left | 195 | -62 | -46 | -10 | 0.000 | 0.160 |
| Precentral Gyrus, Right | 74 | 42 | -8 | 58 | 0.037 | -0.130 |
| Inferior Temporal Gyrus, Right | 68 | 60 | -42 | -12 | 0.043 | 0.140 |
| **Right Lobule V** | | | | | | |
| N/A | | | | | | |
| **Right Lobule VI** | | | | | | |
| N/A | | | | | | |

**Supplementary Table 6***. Differences in cerebellar network connectivity between female reproductive and late postmenopausal groups and relative male control groups, across seeds*. Anatomical regions were identified using the Johns Hopkins University Atlas and the Automated Anatomical Labeling Atlas, Version 3 (Rolls et al., 2020), in MRIcron. Beta values are unstandardized.
